# Benchmark Analysis of Algorithms for Determining and Quantifying Full-length mRNA Splice Forms from RNA-Seq Data

**DOI:** 10.1101/007088

**Authors:** K. Hayer, A. Pizzaro, N. L. Lahens, J. B. Hogenesch, G. R. Grant

## Abstract

The advantages of RNA sequencing (RNA-Seq) suggest it will replace microarrays for highly parallel gene expression analysis. For example, in contrast to arrays, RNA-Seq is expected to be able to provide accurate identification and quantification of full-length transcripts. A number of methods have been developed for this purpose, but short error prone reads makes it a difficult problem in practice. It is essential to determine which algorithms perform best, and where and why they fail. However, there is a dearth of independent and unbiased benchmarking studies of these algorithms. Here we take an approach using both simulated and experimental benchmark data to evaluate their accuracy. We conclude that most methods are inaccurate even using idealized data, and that no is method sufficiently accurate once complicating factors such as polymorphisms, intron signal, sequencing error, and multiple splice forms are present. These results point to the pressing need for further algorithm development.

## Introduction

One of the primary goals of high throughput RNA-Sequencing (RNA-Seq) is to accurately deduce the full-length structure of transcripts. This enables identification and quantification of new splice forms, so that researchers can concentrate their research on the most relevant gene forms in their system of interest. This is a difficult problem however. Most genes are annotated with multiple splice forms and we observe approximately 40% of genes express at least two forms in real data. As methods improve and the number of tissues increases, the apparent complexity will almost certainly go up. Moreover, for species without genome sequence, de novo RNA-Seq and transcript assembly should provide an efficient means of gene discovery in data driven genome annotation.

The nature of current RNA-Seq, resulting in short error prone reads, alignment artifacts, bias introduced by the library construction process, etc., introduces noise into this already difficult problem. Despite this, several algorithms have been developed to infer full-length transcripts *de novo* from short read RNA-Seq data (without using pre-determined annotations), of which Cufflinks [1] is the most widely used. Other methods are Scripture [2], Trinity [3], Oasis (Velvet) [4], Soap-denovo-trans [5], iReckon [6], EBARDenovo [7]. These algorithms can be classified by whether or not they use an alignment to a reference genome sequence. The second approach is to assemble the reads into full transcripts by identifying and assembling overlapping reads. We investigate the accuracy of both types of algorithms below.

A critical but underappreciate aspect of algorithm development is unbiased benchmarking. A few studies investigated algorithms for transcript inference and quantification from RNA-Seq data, the most notable being the RGASP1 (2009) and RGASP2 (2010) competitions [12]. These studies used PCR to validate differences rather than simulated or spike-in data. They concluded that “assembly of complete isoform structures poses a major challenge even when all constituent elements are identified” and that, “Consequently, the complexity of higher eukaryotic genomes imposes severe limitations on transcript recall and splice product discrimination”. They identified that the alignment step that precedes most reconstruction algorithms greatly affects the results so the following year RGASP3 was held to assess aligners [13]. Though a transcript reconstruction paper [12] was published in 2013 from the RGASP effort, it is based on analysis performed in 2009 and 2010 by participants in the first two RGASP competitions.

Between then and now, there has been much new and existing algorithm development, while few papers have analyzed performance. Chandramohan *et al* [14] published an analysis, but the focus was on quantification, not transcript structure. For validation, they used ∼500 genes measured by RT-PCR [15]. However, they relied on early generation public data using single end 36 base reads. Read length has grown (and is still growing) to a standard of approximately 100 bases, and paired-end sequencing has become much more popular. These developments clearly have significant impact(s) on transcript reconstruction and expression quantification. Chandramohan included Cufflinks [1], HT-Seq [16], RSEM [17] and IsoEM [18]. They assessed performance by correlation; however that does not account for the accuracy of absolute intensity levels, just how the values co-vary.

In 2013 the Cufflinks developers released a paper with a fairly extensive benchmarking study using simulated and real data [19]. They generated data with an unpublished in-house method, TuxSeq. The data were generated in idealized fashion similar to one of our data sets, with no polymorphisms and no intron signal. Cufflinks was provided with a perfect alignment sans artifacts. Furthermore, the data was not used to benchmark transcript form reconstruction; instead it was only used for benchmarking differential expression. For that type of benchmarking it is considerably more difficult to generate informative simulated data, as the unknown and extremely complex correlation structure between features would have to be modeled to be able to translate the results of differential expression analysis meaningfully to real data. This, however, is not the purpose of our study. Furthermore the Cufflinks’ study only evaluated Cufflinks’ performance and did not do a comparative analysis to other methods.

Here we update benchmarking of transcript assembly algorithms by using latest generation methods, using synthetic and control datasets. We find that while new methods work well with idealized data and relatively few transcript forms, they largely fail on complex real data. These results point to the need for new algorithm development to solve this difficult, but important, problem.

## Results

Transcript assembly is a difficult problem. For single exons, this is straightforward, one can simply count the tags that overlap and have consistent splice junctions with the exon. Likewise, it is straightforward to determine whether proximal exons are part of the same transcript, as single reads can be mapped across exon/exon junctions with reasonably low error rates. Further, with paired-end data, if the forward and reverse reads are in different exons, then necessarily the two exons are in the same transcript. However, these simple cases don’t apply to full transcript analysis. Transcripts average nearly 2kb and can get considerably larger, and 35% of transcripts have ten or more exons (Figure 1), with the typical gene having multiple splice forms. For example, in mouse suprachiasmatic nuclei of the hypothalamus (SCN), 30% of genes express two or more forms, and 20% express three or more (Figure 2). Sequence polymorphisms, intron signal, and both sequencing artifacts (e.g. error, linker sequences) and artifacts further complicate assembly and quantification of full-length transcripts.

**Figure 1:**
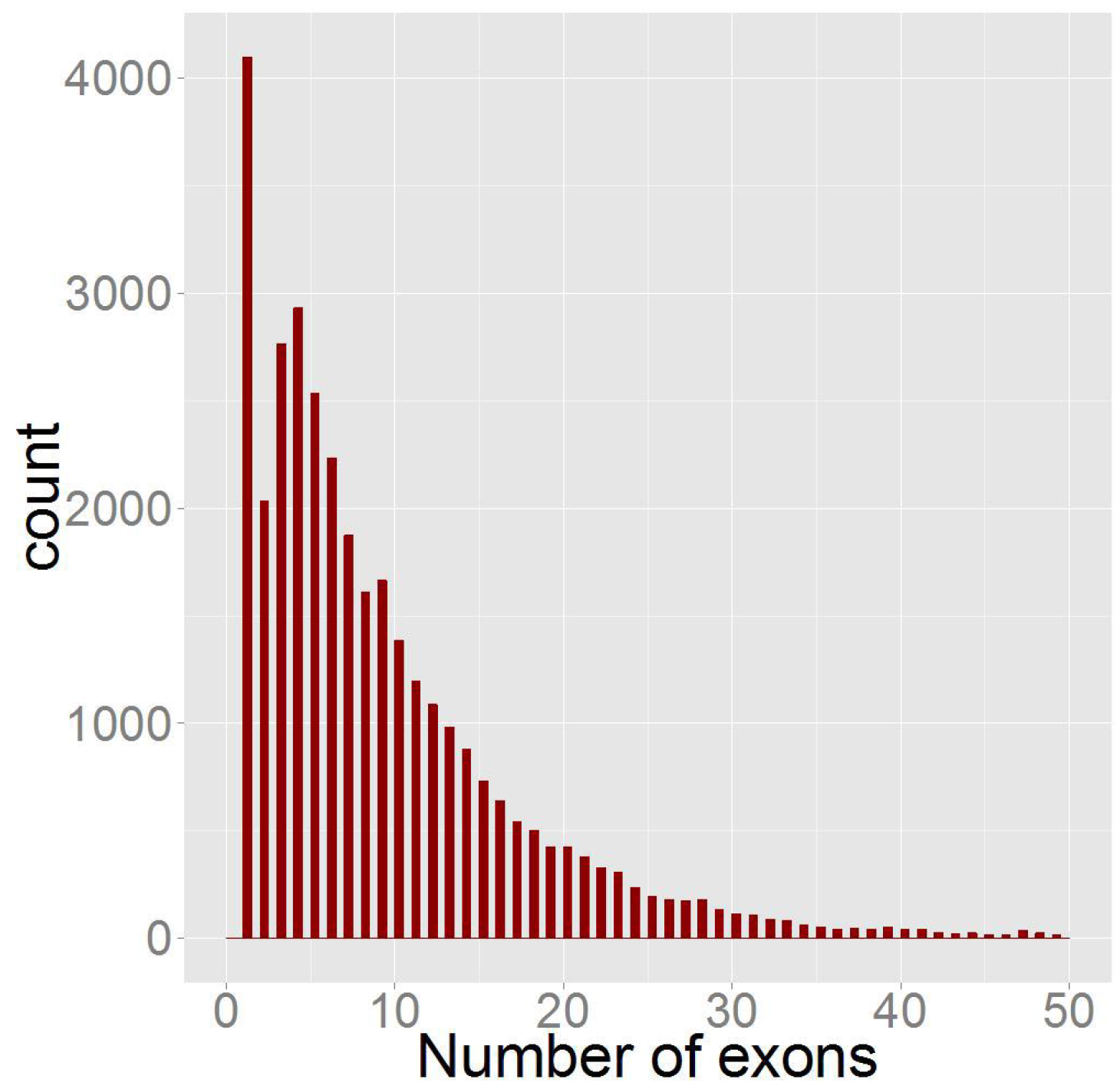
Shows number of transcripts as a function of the number of exons. 90% of transcripts have multiple exons. 65% have more than five and 35% have more than ten.

**Figure 2:**
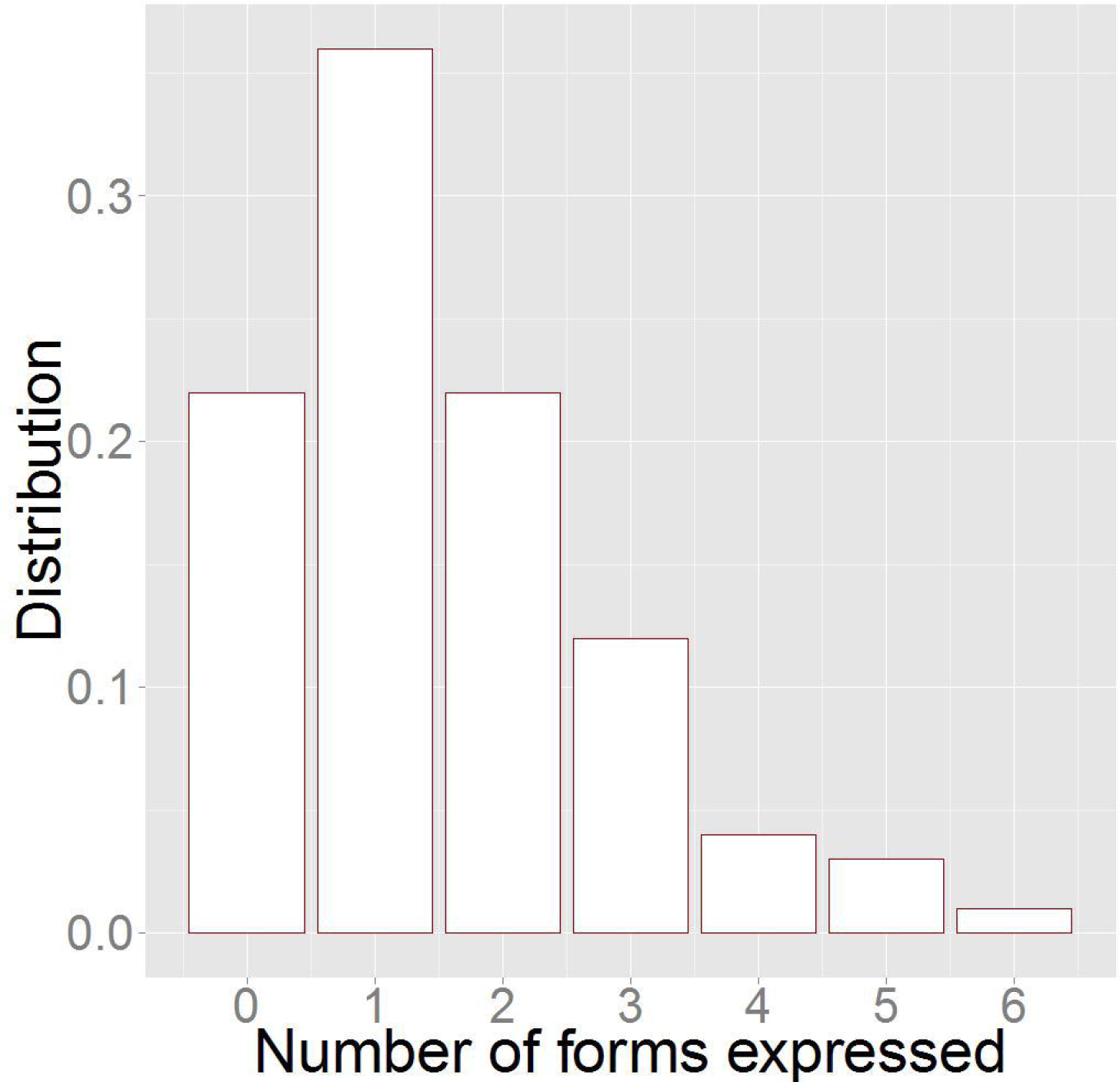
Distribution of the minimum number of splice forms necessary to explain the RNA-Seq junctions in 300M read pairs of mouse SCN. This is based on the first 200 RefSeq genes annotated on Chromosome 1.

Benchmarking is a critical aspect of algorithm development. While it is often included, it is often biased to the advantage of the algorithm developers. Because both the data and algorithms for transcript assembly have changed in the last few years, there is a need for up-to-date benchmarking. We use both simulated and spike-in data to benchmark these algorithms. The advantage of simulated data is that we know exactly where it aligns. The challenge is to simulate data realistic enough for meaningful benchmarking. We used the BEERS simulation engine to generate the simulated data [8]. BEERS starts with a genome and a set of gene annotations; here we used mouse genome build mm9 [9] and RefSeq genes [10], as aligned to the reference genome by the UCSC genome browser [11]. Thus, BEERS does not simulate genomes or gene models, it uses real gene models and simulates RNA-Seq data from those models. We first generated idealized data to determine upper bounds on the accuracy of the algorithms. With idealized data, we minimize complicating factors such as alignment artifacts, polymorphisms, intronic signal, and low expression. If the accuracy is low on ideal data, it is almost certainly worse on real data. We then add the aforementioned complicating factors to simulated data to better mimic real data. With BEERS, we take fragmentation into account yielding data that differ from strictly uniform by the randomness of the fragmentation process.

Spike-in data has advantages of being able to introduce polymorphisms, multiple splice forms, etc., which can all be introduced into cDNAs, and then sequenced with standard methods, thereby capturing all normal sources of techincal error. Because we generated the transcripts synthetically, we know the true form(s) that should be determined by various algorithms if they are working correctly. Importantly, because *in vitro* transcription (IVT) is efficient, the expression of each base pair is theoretically the same. We used 1062 human full-length cDNAs and did IVT-seq. As with simulated data, the full-length transcript forms are known to the base pair. In this data set 50 genes had 2 or more splice forms. These samples were sequenced with poly A-seq and total RNA-seq, the most common protocols, which introduce within-transcript variance (Figure 3) that cannot easily be simulated.

**Figure 3:**
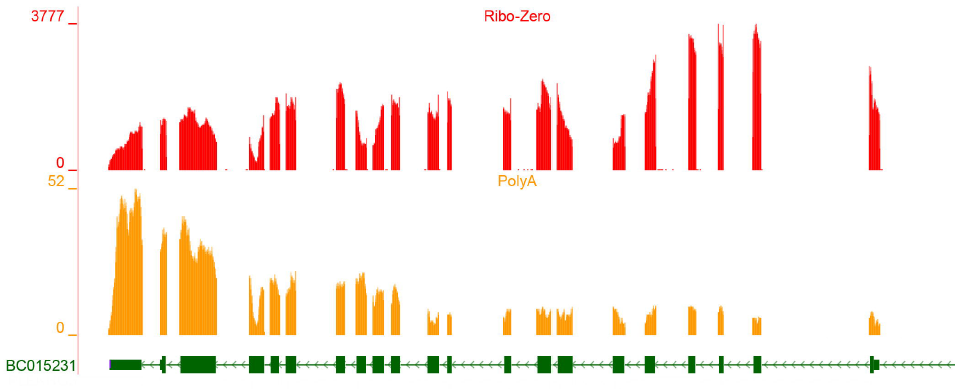
This shows the depth of coverage of a full-length cDNA clone which has been transcribed and subjected to the Ribo-Zero (red) and PolyA selection (orange) protocols for removal of ribosomal RNA. Both protocols result in extreme local bias.

By visually inspecting data in a genome browser using depth-of-coverage plots and representations of spliced reads, it is easy to see that a majority of genes are tractable and the true splice forms can be determined even by hand (e.g. Figure 4). At the other extreme, there are typically a number of regions of extreme confusion observed in real data. Figure 5 shows an extreme number of spliced reads within a gene, and Figure 6 shows an extreme number of spliced reads within one UTR. A handful of such genes typically occur in almost every RNA-Seq experiment and present a major challenge to splice-form level analysis. Another challenge comes from the existence of spliced reads connecting what are annotated as separate genes. Figure 7 shows such an example.

**Figure 4:**
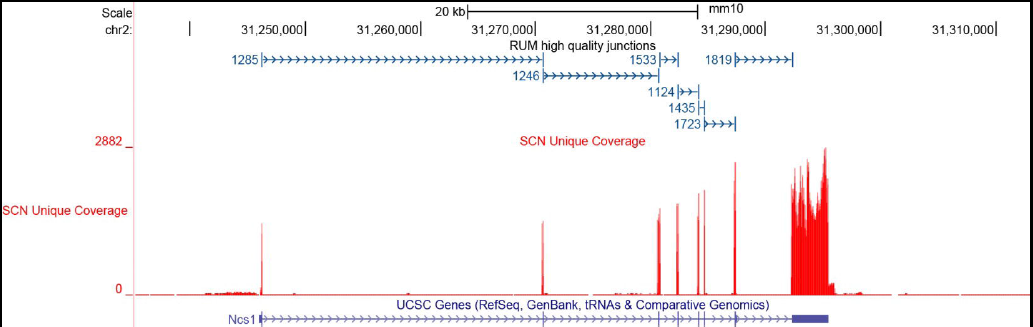
Shows RNA-Seq data for a gene with one (annotated) splice form. The depth-of-coverage is shown in red. Spliced junctions are indicated by the seven directed spans shown in blue. The numbers next to each junction indicate the number of reads that spliced (cleanly) across the junction. In this case there is clear evidence for expression of the annotated form and no evidence for other forms. Note the low level of intronic signal across the entire transcript and extending well beyond it.

**Figure 5:**
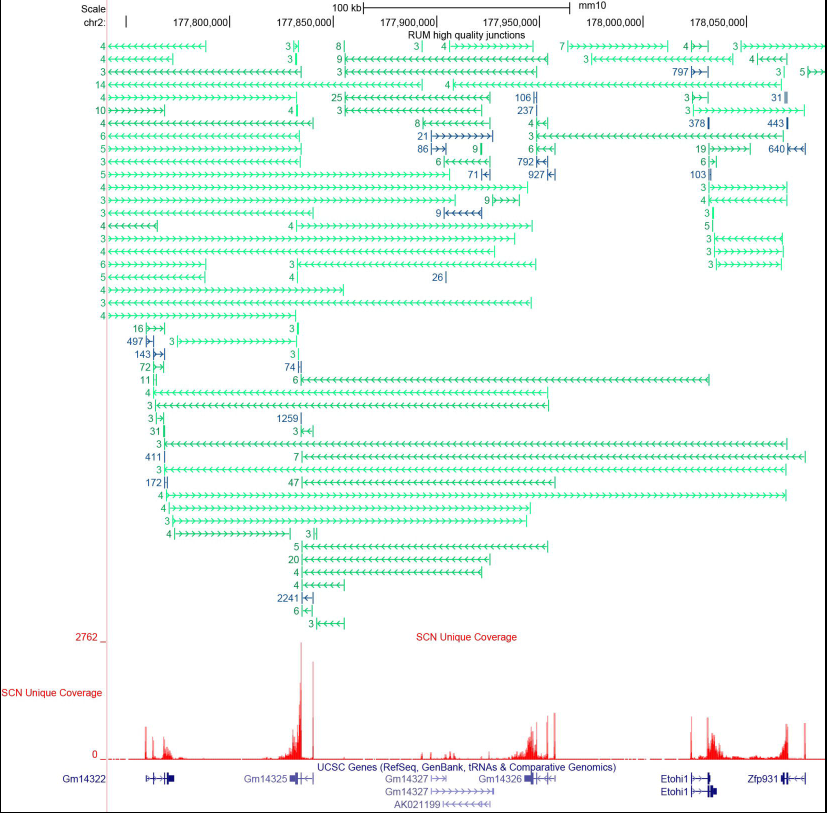
A region of extreme confusion in terms of spliced reads. Junctions are indicated as described in Figure 4, with annotated junctions colored blue and unannotated junctions colored green. These types of genes are seen often in real data and present an extreme challenge to transcript level analysis.

**Figure 6:**
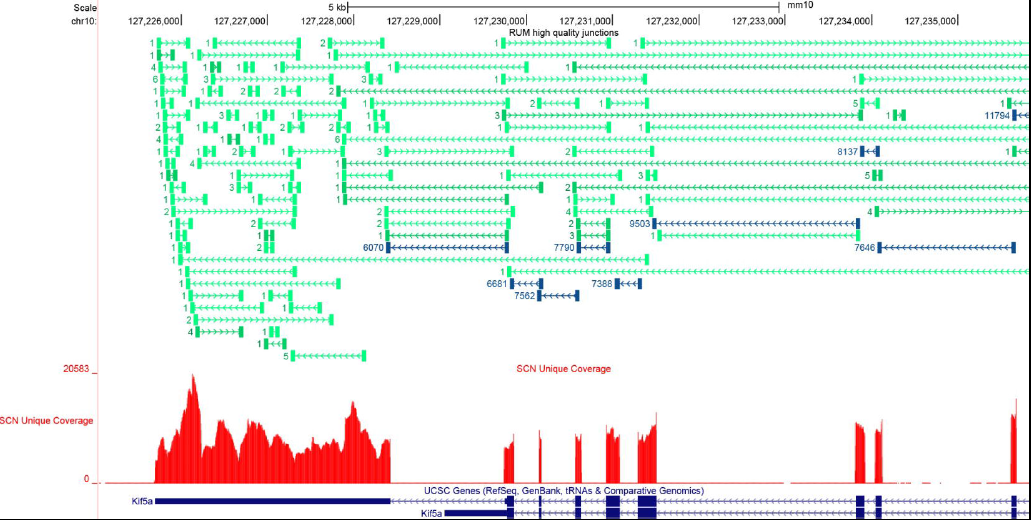
A region of extreme confusion, particularly in the UTR. Junctions are indicated as in Figure 5. Data like this are common in practice, typically observed one or more times per chromosome.

**Figure 7:**
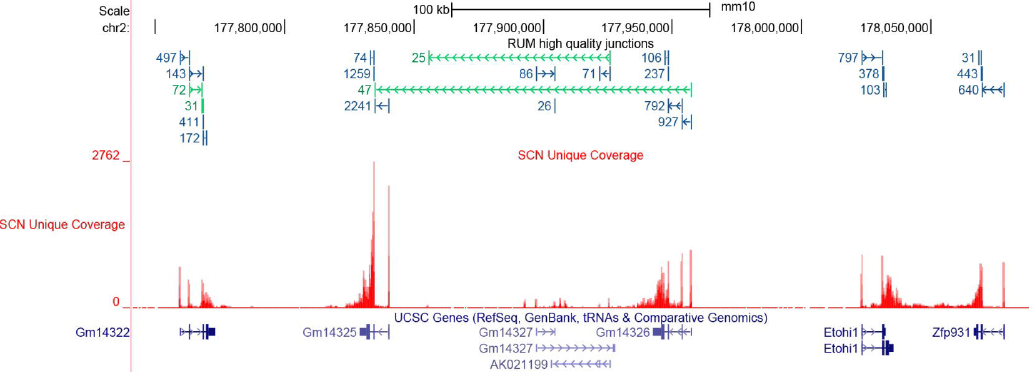
This genome browser track shows data for which reads splice across proximal genes. Junctions are indicated as described in Figure 5. This common and typically unannotated behavior observed in real data present a challenge to transcript level analysis.

### The benchmarking data

We used BEERS [8] to generate two simulated data sets. One was built without polymorphisms, sequencing error, or intronic expression, and with all transcripts highly expressed. In this data, our goal was to assess performance in the easiest scenario. Our Systematic Dataset (SD) has 10,000 genes with numbers of splice forms varying from one to ten. In particular, there are 1000 genes with *n* splice forms, for each *n* = 1,2,…,10, for a total of 55,000 splice forms. There are no overlapping genes and alternate forms for a given gene share both terminal exons, differing only by the presence of absence of one or more internal exons. All genes correspond to RefSeq gene models or are modified from a RefSeq model by exclusion of some number of internal exons. SD has 50 million paired-end reads of length 100 bases per end, with a minimum fragment length of 100 bases, median of 200 bases and max of 400. All genes were expressed at the same depth of coverage of approximately 40 reads.

We also built a simulated Realistic Dataset (RD), with complicating factors similar to what is observed in real, but relatively clean, data sets. In particular this data has a substitution rate of 0.001, an indel rate of 0.0005, an error rate of 0.05, and approximately 30% of the reads coming from introns. The expression levels also follow a spectrùm representative of real data. RD was generated with the full and unmodified RefSeq annotation track of 30,150 transcripts, again with 50 million paired-end reads of 100 bases per end and with a minimum fragment length of 100 bases, median of 200 bases and max of 400.

Finally, we used data from *in vitro* transcription of 1062 human cDNAs (IVT), sequenced by both poly A seq and total RNA-seq. Because these are cDNAs, we know the exact nucleotide sequence, including all the exòn-exon junctions. Of these genes 50 have multiple splice forms. This data provides an evaluation of each algorithm on real data.

Some algorithms (e.g. Cufflinks, CEM, and iReckon) have a mode where they take a set of annotated transcripts as input and then determine which forms are expressed. To benchmark these methods properly, we needed to augment SD with a set of transcripts that are not expressed. Therefore, we generated an additional set of 55,000 splice forms that are not expressed, by selectively removing exons. We then gave the algorithms the annotation file with all 110,000 transcripts, half of which are expressed at a moderate level and half are not expressed at all. These data sets with these simplified annotation tracks represent the easiest case of perfect data, so their error rates should provide an estimate of a lower bound of performance in practice. Results are considered as functions of the number of splice forms. In real data, we observed roughly 45% of expressed genes express one form, with roughly 10% expressing more than four forms (Figure 2).

For alignment-guided methods, we aligned the reads with TopHat2 [20] (hereafter referred to as TopHat) and STAR [21]. Prior studies of alignment algorithms found STAR to be a close second to the most accurate aligner (GSNAP [22]), however STAR runs orders of magnitude faster than any other RNA-Seq alignment algorithm. Because of its accuracy and rising popularity, we used STAR to represent the high end of aligners. TopHat was included because it is the most widely used and because it is the recommended input for Cufflinks. For the simulated data, we also have perfect alignments with no alignment artifacts. The alignment-guided algorithms produce gene models consisting of sequences of connected exons given by genome coordinates. It is on this level that the results of all such algorithms become directly comparable.

### Accuracy metrics

We compared the putative transcript models produced by each algorithm with the known models used to generate the data to calculate an accuracy metric. If a transcript model agrees with a known model on all *internal* splice junctions, we call it a true positive. As the start and end of transcription are notoriously difficult to predict, we do not consider them in our benchmarking. If at least one of the internal junctions is wrong, we call it a false positive. If a model is expressed but not reported by the algorithm, we call it a false negative.

When predicting transcript structure as a sequence of exon start and end sites, there are an astronomical number of false negatives – any sequence of an even number of consecutive coordinates is a putative transcript. Therefore, as long as the algorithm itself does not return an astronomical number of putative transcripts, the false positive rate will be very close to zero and the specificity will be very close to one. This remains true even if every predicted transcript is a false positive. Instead it is much more informative to report the percent of predictions that are false positives, known as the False Discovery Rate (FDR) [23]. We also report the False Negative rate (FNR), i.e. the number of false negatives as a percent of all expressed transcripts.

### Performance of the Algorithms

We evaluated algorithms in two transcript inference problems. The first was to determine which transcripts are expressed from a given set of annotated transcript models. The second was to determine *de novo* the forms of the expressed transcripts, which is a considerably harder problem.

While most algorithms perform well with perfect data and a single splice form, they tend to falter when predicting multiple splice forms. Figure 8 gives the performance on SD in terms of the FDR and the FNR, for the various algorithms. Results are given separately for genes with one splice form and genes with two splice forms. Not surprisingly, several algorithms (e.g. Cufflinks, CEM, iReckon) perform quite well when trying to infer the expression of a gene with one splice form, when provided with accurate annotation and a perfect alignment. Once there are two splice forms, all algorithms have a > 10% FDR. The application of Cufflinks using a TopHat alignment (Cufflinks+TopHat), which is common in practice, results in a 40% FDR. On real data the FDR must be even higher. Not surprisingly, the *de novo* methods incur substantially higher error rates, with Trinity having a nearly 90% FDR on two-splice-form genes, coupled with an approximately 50% false negative rate. Curiously Cufflinks performs better with a TopHat alignment than with a STAR alignment, even though STAR produced a more accurate alignment.

**Figure 8:**
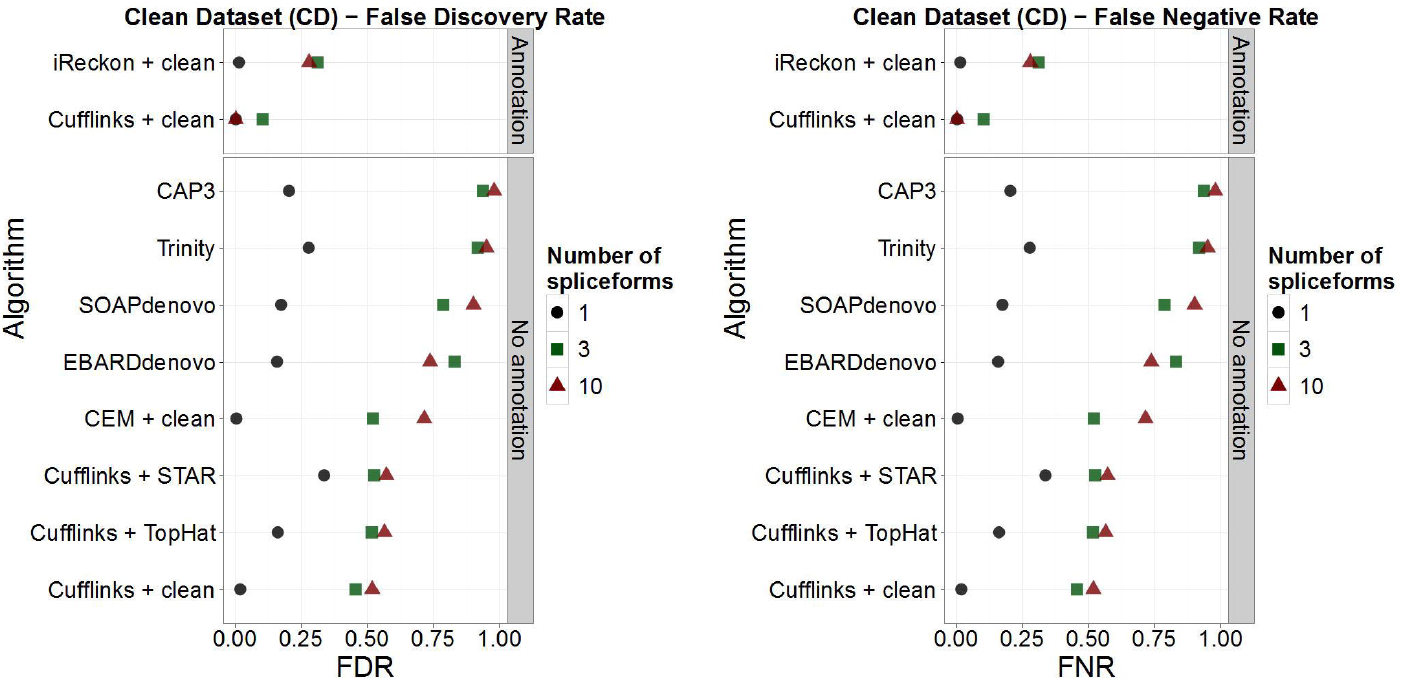
False Discovery Rate (FDR) and False Negative Rate (FNR) for CD. Graphs show results for genes with one three and ten splice forms, each category of which is represented by 1000 genes.

Because of its wide use in practice we chose Cufflinks as an exemplar and examined its performance with increasing numbers of splice forms. Figure 9 breaks down for SD the FDR, FNR and True Positive (TP) Rate for Cufflinks plus TopHat, separated by number of splice forms per gene ranging from one to ten forms. With one or two forms, Cufflinks performs satisfactorily. However, as three or more forms are considered, the FDR rises and the true positive rate falls precipitously. To get an idea of what is causing the false positives we inspected specific examples produced by Cufflinks (see example in Figure 10). Usually, Cufflinks correctly identified the local characteristics of a transcript but failed in correctly predicting the full-length transcript. In many cases, split transcripts were produced. Other algorithms also had this problem.

**Figure 9:**
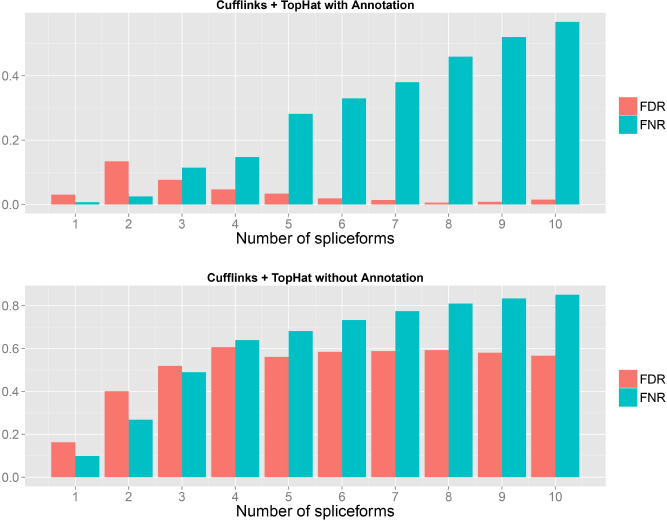
Breakdown for CD of the FDR and FNR for Cufflinks inferred genes, with up to ten splice forms. Data were aligned with TopHat2.

**Figure 10:**
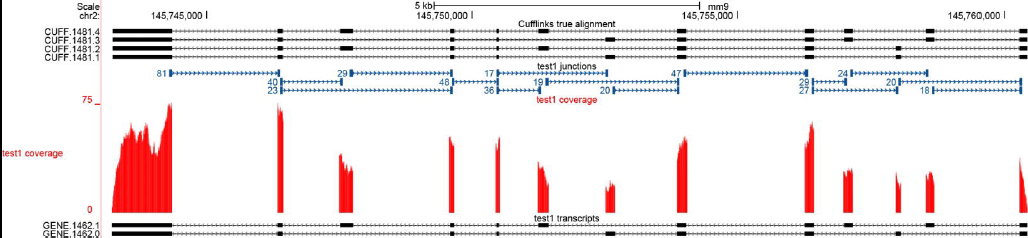
This genome browser graphic shows four transcripts returned by Cufflinks (top) on test CD100, as compared to the two real transcripts the data were generated from. The red track shows the depth of coverage plot for the true alignment. Local alternative splicing events were joined in non-existent ways to create the two false positives.

RD represents more realistic data, with polymorphisms (in the form of substitutions and indels), sequence error, and intron signal. Figure 11 shows the results for genes with two splice forms. For this data set we expanded the comparison to include other aligners and combinations. Cufflinks performs adequately if it is given a true alignment and accurate annotation. The application of Cufflinks + TopHat to perform *de novo* identification incurs an FDR error rate around 30% with FNR around 25%. This is slightly better than what we observed with ideal data (Figure 8), which is because there are many more splice forms in the SD set (e.g. 1000 genes with 10 forms each), while the RD data is comprised of Refseq genes with fewer forms.

**Figure 11:**
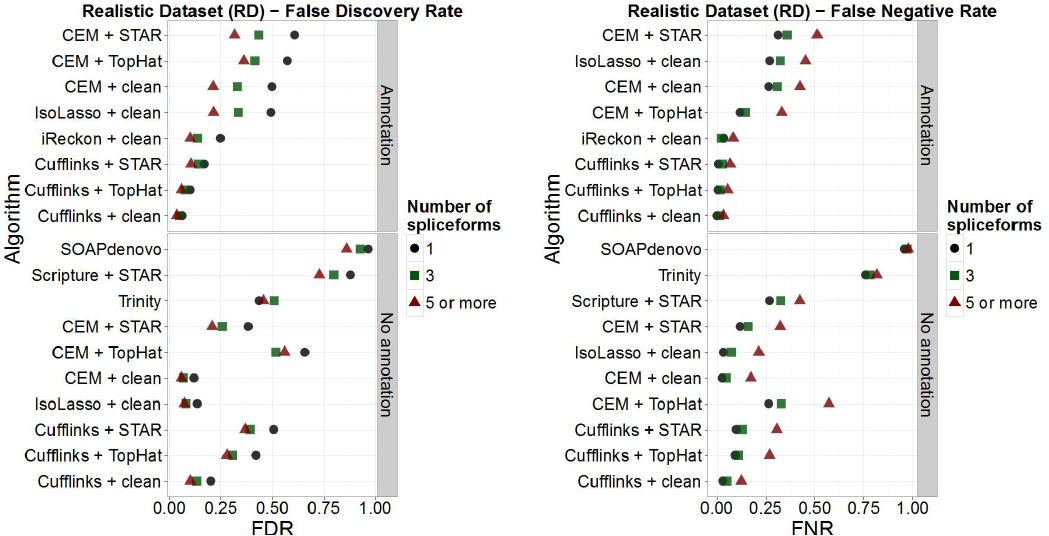
Accuracy results for RD for genes with one, three and five or more splice forms.

We expect the results for RD are still quite conservative and that true sequencing artifacts would introduce considerably more complexity. This is demonstrated by the results on the IVT data (Figure 12). In this case, since this is real sequence data, we could not provide an error-free alignment. The annotation given was of the known structure of the 1062 transcripts. With this data, both the FDR and FNR were much higher. Cufflinks + TopHat + Annotation performed the best, with a FDR of 0.2 and a FNR of 0.26. This was an order of magnitude worse than for the SD data, which speaks to the complications introduced by alignment errors, polymorphisms, etc. For the genes with more than one splice form, the results were worse still.

**Figure 12:**
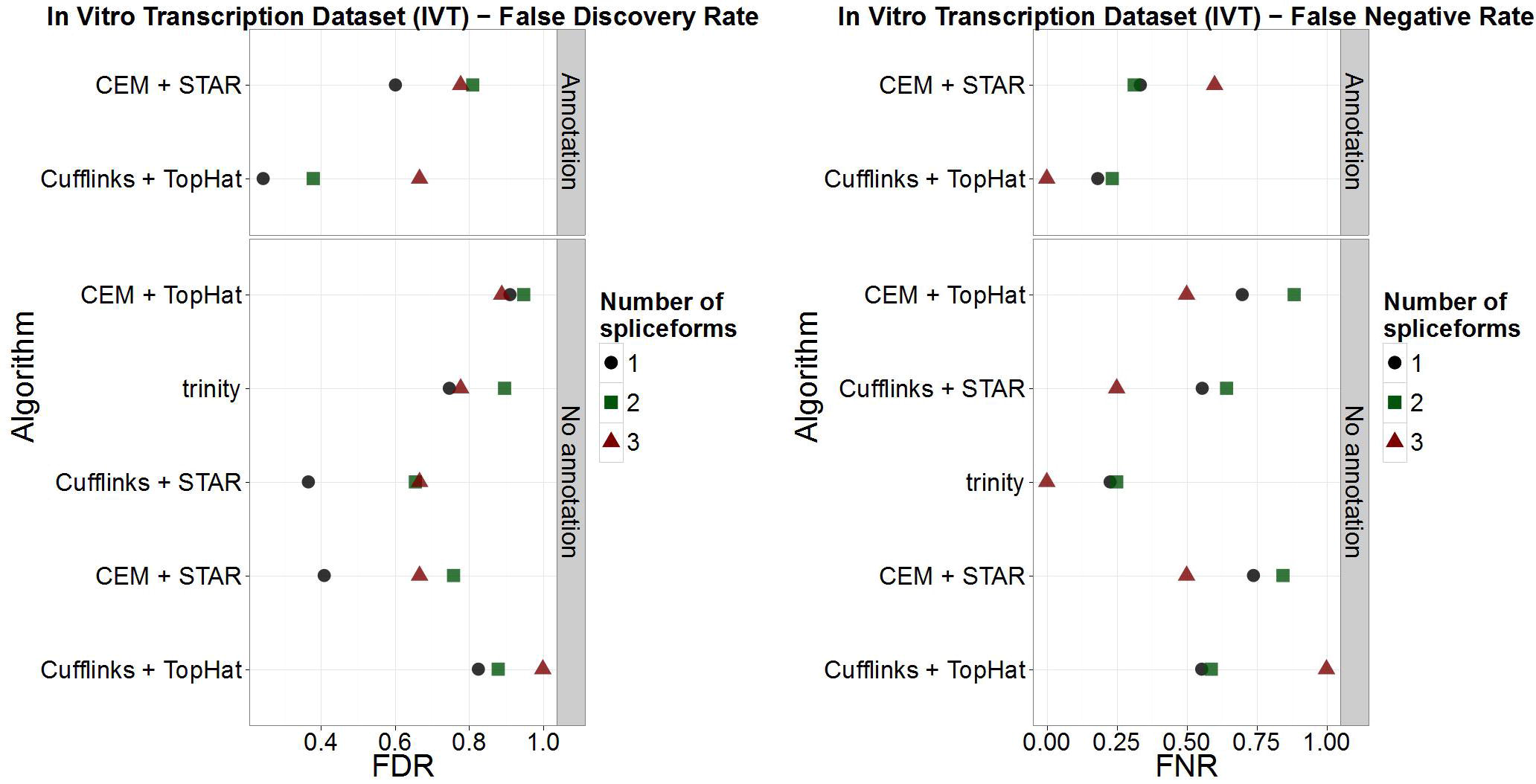
Accuracy results for the spike-in data set for one, two and three transcripts per gene.

We now turn to expression estimation. This can only be assessed by RD since the others are not quantitative (all transcripts are expressed equally). Since there is no “true” expression value for incorrectly inferred splice forms, it only makes sense to assess the quantification of known annotated features. Figure 13 shows the scatter plot of Cufflinks’ inferred FPKM values versus the true FPKM values. This represents the ideal case where Cufflinks was run with the perfect alignment and annotation. As accuracy metric we calculate the percent of transcripts where the FPKM was more than one logarithm from the truth. In this idealized data the metric is 16.76%. The points on the *y*-axis are transcripts that have true FPKM equal to zero but which were given positive FPKM by Cufflinks, while on the *x*-axis are transcripts that have positive true FPKM but were given an FPKM of zero by Cufflinks. If Cufflinks is not provided with annotation this number rises to 38.62%. If Cufflinks is run with TopHat and without annotation is rises to 54.12% (Figure 14). Figure 15 shows the performance of the other relevant algorithms. These results are not encouraging.

**Figure 13:**
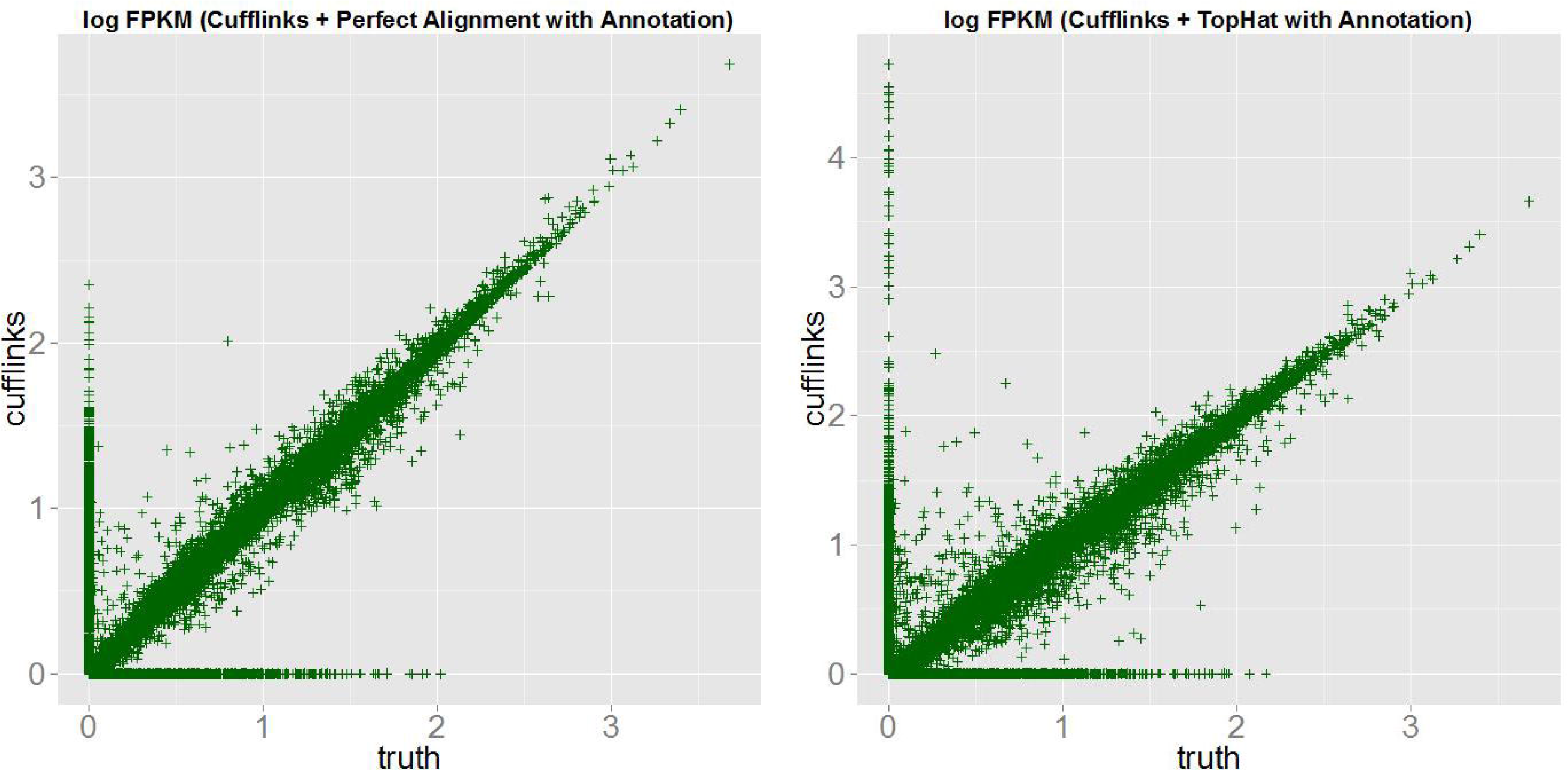
Scatter plot in log scale comparing the true FPKM values to the Cufflinks inferred FPKM values on RD using the perfect alignment and with Cufflinks given the true annotation. This represents the most ideal case. 16.76% of FPKM values differ by more than one log from the true value. Replacing the perfect alignment with the TopHat alignment increases this number to 24.65%.

**Figure 14:**
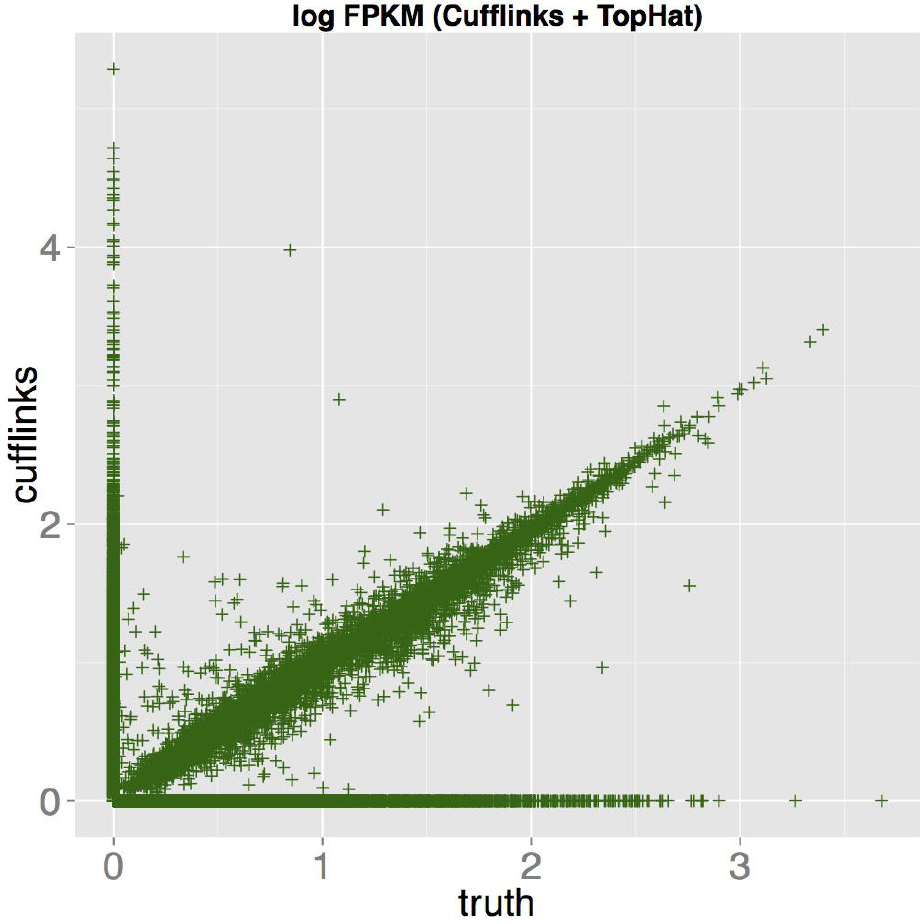
Scatter plot in log scale comparing the true FPKM values to the Cufflinks inferred FPKM values on RD using the TopHat alignment and without annotation. This represents a common use case in practice. 54.12% of FPKM values differ by more than one log from the true value.

**Figure 15:**
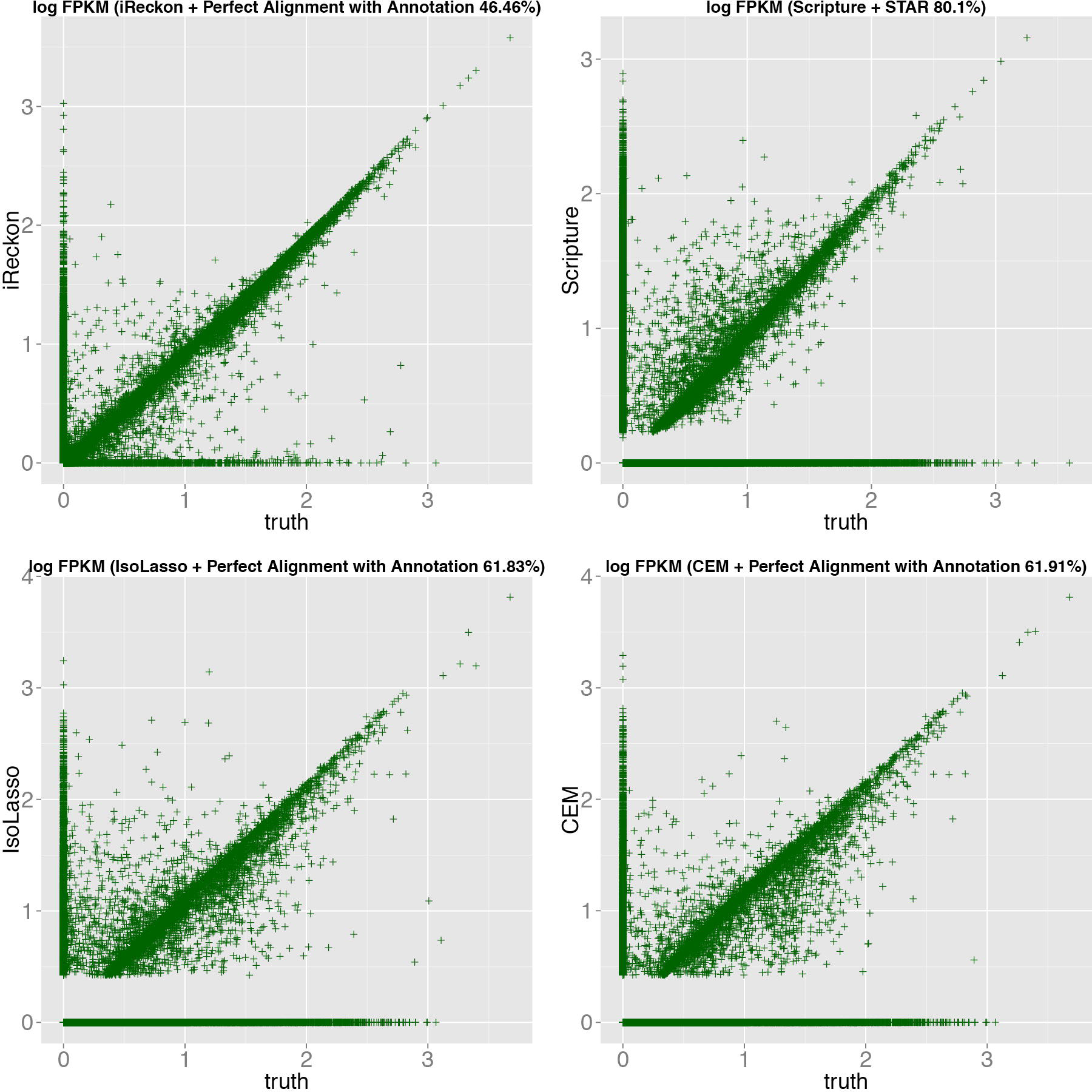
FPKM scatter plots in log scale comparing inferred FPKM values to the true FPKM on RD for various algorithms. The percents give the percent of transcripts that are more than one log off from the true value.

## Discussion

Transcript structure determination, either *de novo* or with annotation, is a challenging problem. To see how latest generation algorithms do, we devised a set of tests using simulated and *in vitro* transcribed RNA-Seq data. We generated one clean data set (SD) simulated from 10,000 genes with 1-10 splice forms. SD used 100 bp paired end sequencing. As this is clean data, it had perfect representation of all bases (save the first and last exons), no sequencing or mapping errors, and no polymorphisms. We also simulated a 100 bp paired end realistic data set (RD) that from ∼30,000 Refseq transcripts. We generated this data set with polymorphisms, error rates, and uneven representations of internal exons, at rates consistent with what we and others observe with high quality data in practice. Finally, we did *in vitro* transcription on 1062 full length human cDNAs, where mapping errors, uneven base representation, and polymorphisms are unavoidable. With all of these data sets, we know the truth, making them ideal for evaluating transcript assembly algorithms.

On *perfect* data (SD) with a *single* splice form (Fig. 4), Cufflinks, CEM, and iReckon all do reasonably well, with FDRs < 0.1. However, while Cufflinks has a low FNR with this ideal data, CEM, iReckon, Ebard, SOAP, Cap3, and Trinity do poorly (FNRs from 0.3 to 0.9). With clean data and just two forms per gene, FDRs for all algorithms go up considerably. Cufflinks does the best, with a FDR of 0.16, with the rest of the algorithms ranging from 0.3 (iReckon) to 0.85 (Cap3 and Trinity). Put simply: algorithms designed to delineate transcript forms make many false discoveries, even on perfect data. Save Cufflinks, false negative rates for all algorithms are also high.

Cufflinks, which performed the best with this data, did considerably worse (minimum FDR of 0.5) when there were three or more transcripts per gene, which is arguably where transcript assembly algorithms are needed most (Fig. 5). The other algorithms did even worse on this data..

With the RD data, we sought to model in realistic errors, e.g. mapping errors, polymorphisms, intron retention. RefSeq transcripts are simpler, on average, than those that were used in SD, with fewer alternate forms differing by multiple exons. Therefore some of the reported error rates are lower in RD because there are more genes on average with simple alternate splicing than is represented in SD. RefSeq, however, is significantly incomplete with much more complex splicing observed in real data than is reflected in RefSeq annotation. Therefore the error rates in SD are specific to genes with a particular structure of alternate splicing and should not be directly compared to RD without breaking down the genes into more specific categories than just the number of transcripts.

The IVT data (IVT) is perhaps the most informative control dataset, because rather than modeling errors, alignment, polymorphisms, base representation, it simply has them as they occur naturally. FDRs with this data are considerably higher, even though 95% of genes had only a single form. Tophat and Trinity had FDRs of about 0.2, with the others ranging from 0.33 to 0.92. FNRs were likewise high, ranging from 0.27 (Cufflinks + annotation + RUM or Tophat) to 0.87 (CEM + RUM or Tophat). Without annotations,which is appropriate for detection of unannotated transcripts, FDRs ranged from 0.5 to 0.9 and FNRs ranged from 0.4 to 0.87. Put simply, on this real, benchmark data, with 95% of transcripts having only a single form, there were as many false transcripts predicted as true ones, and half of the true ones were missed. In the IVT data, we had a limited subset of ∼50 genes with more than one transcript. On this limited set of real data, performance was still worse.

It is instructive to look at what aspects of the popular algorithms need improvement in order to increase their accuracy. Cufflinks uses junction and coverage information to determine local alternative splicing events. But it is willing to assemble the local information into global transcripts in all possible ways that are consistent with the short reads, and this is the source of many of the false positives. Since the integrity of the transcripts is lost in this approach it is not clear what advantage it could have over an exon/intron/junction level analysis. Another problem is that it does not effectively sort out exon from intron signal. In most RNA-Seq data sets there is some amount of signal at almost every base of almost every intron. This intron signal is often assembled by Cufflinks into long exons, resulting in many more false positives. Perhaps reading-frame information could be used to limit this type of artifact. There is also generally a smattering of junctions that connect exons to the middle of introns yet probably do not represent real exon/exon junctions. The challenge is for algorithms to find a way to ignore the intron and spurious junction signal to get at the clean transcript models.

Scripture uses an even more liberal approach by essentially ignoring junction information and instead taking a “peak finding” approach to identify putative exons and then connecting them in all possible ways. This approach also suffers from the fact that there is insufficient information to connect local inferences into global inferences and so it constructs a large number of false positives by joining local effects to make full-length transcripts. We have also observed that Scripture tends to find many peaks in introns where the signal was generated via a uniform model, and so should not have any significant peaks. The extreme overcalling of forms makes it unclear how to utilize the output of Scripture in a practical way.

The de novo methods such as Trinity are trying to solve a much harder problem by not using the information coming from the genome sequence. If no genome exists then there is not a lot of choice and this approach might provide information that can be informative. However, for model organisms, and many others with a reference genome available with some degree of community annotation, it is hard to imagine any benefit of using a de novo approach.

We have seen in Figures 5-7 regions that present a particular challenge to transcript level analysis. Indeed it is not even clear that these are true splicing events and not just artifacts of sequencing or alignment. So the first task in using RNA-Seq to perform transcript level analysis should be to determine what part of these challenging regions represents true biology that is worth quantifying and what part of it is artifactual.

### Conclusion

This problem is fundamental and virtually all RNA-Seq investigators desire to do their analysis cleanly at the transcript level. However, short reads fundamentally lack the information necessary to build local information into globally accurate transcripts. The strong desire of the community is what has made it possible for several algorithms to gain widespread use in spite of the apparent impossibility of a clean solution to date. This underscores the importance of more research into this problem. Most likely it a satisfactory solution will involve an evolution in the nature of the data. Or perhaps some keen insight into how to identify and effectively utilize signals in the genome that inform cellular machinery on what splice forms to generate

## Methods

### Simulation

The simulator BEERS was used to generate two different datasets of 50 million paired end reads. SD was generated with the following parameters “-readlength 100 -fraglength 250 -error 0 - subfreq 0 -indelfreq 0 -intronfreq 0 -palt 0” with fragment length distribution min/median/max=100/200/400. RD was generated with the following parameters: “-readlength 100 -fraglength 250 -error 0.005 -subfreq 0.001 -indelfreq 0.0005” with fragment length distribution min/median/max=100/200/400.

Simulated data set SD is available here:

http://itmat.greg.s3.amazonaws.com/hayer1_SD.tar.gz

Simulated data set RC100 is available here:

http://itmat.greg.s3.amazonaws.com/hayer1_RD.tar.gz

### Alignments

Reads were mapped using several commonly used mapping tools: TopHat (version 2.0.8b), RUM (version v2.0.5_05) and STAR (version 2.3.0e). The following flags were used for the TopHat alignment: “-read-mismatches 5 --read-edit-dist 5 --read-gap-length 3 --read-realign-edit-dist 0 --mate-inner-dist 0 --max-insertion-length 6 --max-deletion-length 6 --min-intron-length 30 --splice-mismatches 1 --no-discordant --microexon-search”. The following flags had to be added to the STAR aligner to achieve compatibility with Cufflinks: “--outSAMstrandField intronMotif --outFilterIntronMotifs RemoveNoncanonical”. (This will add the XS tag to the SAM file.) The aligner RUM was used with default parameters.

### Genome guided reconstruction algorithms

Genome guided reconstruction algorithms were run with the recommended default parameters. The list of tested algorithms included Cufflinks (version 2.0.2), Scripture (beta version 2.0), CEM (version 2.5.2), IsoLasso (version 2.6), Casper (version 1.1.2) and IReckon (version 1.0.7). Some of the given algorithms can be provided with gene models to improve accuracy. The algorithms were run in both modes if available. Only gene models with sufficient coverage or confidence value were used in computing the false negative rates.

### De-novo reconstruction algorithms

The following de-novo reconstructions algorithms were run on the simulated data sets: Trinity (version r2012-10-05), OASES (version Feb 1, 2010), SOAPdenovo-Trans (version 1.0) and EBARDenovo (version 1.2.2). All tools were run with default settings. To fairly compare the de-novo reconstruction algorithms to the genome guided tools, the obtained transcripts were aligned back to the mm9 reference with GMAP (version 2012-07-20). Finally only transcripts that were assigned successfully were used to evaluate the algorithms performance.

### Performance evaluation

All results were evaluated with Ruby scripts, which are available here: https://github.com/khayer/benchmarking_scripts. For the purposes of this work the scripts not only return the false positives, true positives, etc. but also split the results by splice-forms and FPKM-values (which only makes sense in our case for RD).

### Spike-In Data

In vitro transcription RNA was derived from an amplified plasmid library taken from the Mammalian Gene Collection. The RNA was pooled and prepared with the Ribo-Zero Gold kit (Epicentre catalog no. RZHM11106). Afterwards the RNA was converted into Illumina RNA-seq libraries with the TruSeq RNA sample prep kit (Ilumina catalog no. FC-122-1001) and sequenced with an Illumina HiSeq 2000 (paired 100 bp reads).

